# Identification of Novel Transcripts and Exons by RNA-Seq of Transcriptome in *Durio zibethinus* Murr

**DOI:** 10.1101/2022.10.08.511396

**Authors:** Nurul Arneida Husin

## Abstract

*Durio zibethinus* is a popular seasonal fruit among the Asian population and is frequently associated with economic losses due to a rapid post-harvest process. To identify novel transcripts and characterize *D. zibethinus* transcriptome, we employed high throughput RNA-Seq analysis. In this study, we characterized the growth and development of *D. zibethinus* at different growth stages, including the young, mature, and ripening stages of growth. After mapping, 110 million high-quality reads were analysed for each of the nine samples, revealing that each contains in range of 76,700 to 89,117 transcripts and 561,211 to 646,291 exons. With Cuffcompare to obtain the unannotated transcript, the novel exons, introns and loci were identified in the developmental stage with value of 3,438/313,476 (1.1%), 1,657/245,129 (0.7%) and 1,197/44,509 (2.7%), respectively. The differential expression genes (DEGs) in up-regulation were compared under the three growth stages: 1,496 (YS/MS), 2,154 (YS/RS), and 2,153 (MS/RS). These genes were then subjected to clustering analysis. A comprehensive study from the heatmap indicates that response to stress, hormone signalling, cuticle formation, general transporter, receptor and protein kinase and transcription factors are highly regulated gene functions for all growth stages. In the present analysis, we emphasized identifying and classifying unknown important transcripts, disregarding transcripts with accurate annotations. Differential expression analysis identified 280 unknown significant transcripts, presumably involved in various biological functions in durian growth. Top blastn results for 183 unknown transcripts revealed homology (80%–100%) to Bombax ceiba, Gossypium hirsutum, Gossypium raimondii, Herrania umbratica, Juglans regia, and Theobroma cacao. Under the category of unknown transcripts without a Blastn match, two unknown transcripts match GO terms. TCONS 00034019 and TCONS 00058246 match the InterPro GO Names for Biological Process (P) P:GO:0046622; P: positive regulation of organ growth. In the category of sequences with no BLASTx hit and no IPS match, three ORF sequences longer than 300 nt were identified. Our results significantly improved D. *zibethinus* transcript annotation and provide valuable resources for functional genomics and studies in durian.

## Introduction

Genomic research of durian is still at a nascent level, and it needs to be accelerated. An in-depth understanding of Durian fruit molecular biology can help to mitigate its strong offensive aroma through gene regulation. In addition, the prolongation of the short post-harvest life of the fruit may be considered for further genetic improvement of the Durian (Husin et *al*, 2018). The strong and pungent durian scent is caused by sulfur-containing compounds, while the fruity smell is caused by esters and alcohol (Siriphanich, 2011). Combining positive traits, fewer odours, and high-stress tolerance is a potential way to grow superior Durian clones in the future.

High throughput sequencing technologies are routinely used for global transcriptome profiling at an unprecedented scale due to their ability to provide detailed information on transcript abundances. Several scientific studies have suggested that deep sequencing is possible to detect differential expression of genes, novel genes and transcripts differentially expressed isoforms and splice variation unique for each condition (Jakhesara et *al*., 2013). Although the majority of the genome is translated into RNA, only few of these RNAs are translated into proteins. Noncoding RNAs (nRNAs) are the name given to these RNAs that do not develop into proteins. Although ncRNAs have been known for some time, their inadequate annotation makes RNA-seq data difficult to interpret (Weirick et al., 2016).

While the durian genome and transcriptome (Musang King var.) has been published, no prior knowledge was generated about the novel and unknown genes or transcripts expressed in the durian transcriptome. The final reference assembly consisted of chromosome-scale pseudomolecules with a length of 30 pseudomolecules larger than 10 Mb and comprising 95% of the 712-Mb assembly (the pseudomolecules are hereafter referred to as chromosomes, numbered by scale). These numbers closely correspond to previous estimates of the durian number of haploid chromosomes (1n=28, 2n=56) (Teh et *al., 2017*).

Transcriptome data and associated annotation data may fill in some gaps in Durian molecular biology, which is often focused on phytochemical compositions and nutritional properties. This research is expected to result in the discovery of new transcripts and exons that will aid in the genetic enhancement of durian. The results of transcriptomics research can be used to investigate new and unknown genes for potential and commercial uses. The new data set generated by high throughput sequencing technology will be helpful in genomes and transcriptomics research on the Durian plant. This paper reports the identification and characterization of novel transcript and exons from the transcriptome of Durian (D24 var.) fruit pulp in its three stages of development (young, mature and ripening). We present several novel transcripts expressed in *D. zibethinus* (D24 var.).

## Material and Methods

### Plant Material

Nine biological replicates of the Durian variety (clone D24) were freshly collected at Agriculture Land UPM at the young, mature, and ripe stages. Fruit pulp tissues were separated from the husk (and seeds), frozen in liquid nitrogen, and stored at -80 degrees Celsius until complete RNA isolation. Young fruits were harvested at the age of 90 days. The mature fruit (120 days) was left to ripen naturally at room temperature for 7 days (for a total of 127 days) to obtain pulp tissue samples during the ripening stage.

### RNA isolation, cDNA library construction and HiSeq Illumina sequencing

The RNA extraction kit was used to separate the RNA, named the GeneAll RibospinTM Seed/Fruit RNA mini kit (GENEALL BIOTECHNOLOGY CO., LTD). Using the NEB Next Poly(A) mRNA isolation module, poly-A mRNAs were purified from 1 µg of total RNA. The first and second strands of cDNA synthesis were conducted in the NEB Next RNA Library Prep kit with purified mRNA (Illumina). All protocols were conducted on the manufacturer’s instructions. Before sending it for sequencing, the library’s consistency and volume were measured by the Qubit 2.0 Fluorometer and Agilent Tape Station. Nine pre-prepared cDNA libraries have been submitted to the NGS service provider (Novogene, China). The libraries were sequenced using a 2 × 150 bp pair-end protocol with HiSeq Illumina Platform. The data output predicted was 16.7 million reads or 5 GB per sample. The total data for the 9 samples was 150.3 M, or 45 GB.

### Pre-processing of Illumina RNA-Seq reads using FastQC and Trimmomatic

FastQC is a Java software used to check the quality of the transcriptome. The purpose of running FastQC is to determine the quality encoding of fastq files, to assess the quality of sequencing samples, and to identify overrepresented sequences-adapters and other potential contamination that may occur in the sequencing process. FastQC was run to check the quality of the raw data and the quality of the trimmed data. The comparison was made before and after (twice for each sample) to ensure only the cleaned reads are used for downstream analysis (Andrews, 2010). Trimmomatic is a tool that performs a variety of trimming of the Illumina FastQC of paired-end or single-end data. It removes the added adapters during the sequencing process (Blankenberg et *al*., 2010). The parameter settings in trimmomatics are as follows; PE= Paired-end (two separate files), ILLUMINACLIP= true, standard adapter, max mismatch=2, SLIDINGWINDOW= number of bases to average across= 1, average quality required=20, HEADCROP= 15, MINLEN= 100, LEADING=3, and TRAILING=3.

### Heatmap Sample Clustering

To explore the similarity of the nine samples, the samples-level QC was applied using hierarchical clustering methods. The distance function was applied to the transpose of the transformed count matrix to get the sample-to-sample distances. Heat maps were constructed for all three groups to access similarities and dissimilarities between samples using the transformed data. The sample level QC was observed to see how well the replicates clustered among themselves. Also, the test reveals if any sample outliers should be removed before analyzing DE. Figure 1A shows the gene expression correlation between young and mature levels. The hierarchical tree indicates that the clustered samples are similar. The dark blue colour blocks represent the substructure in the mature stage sample, replicating data, and high similarities.

**Figure 1.**
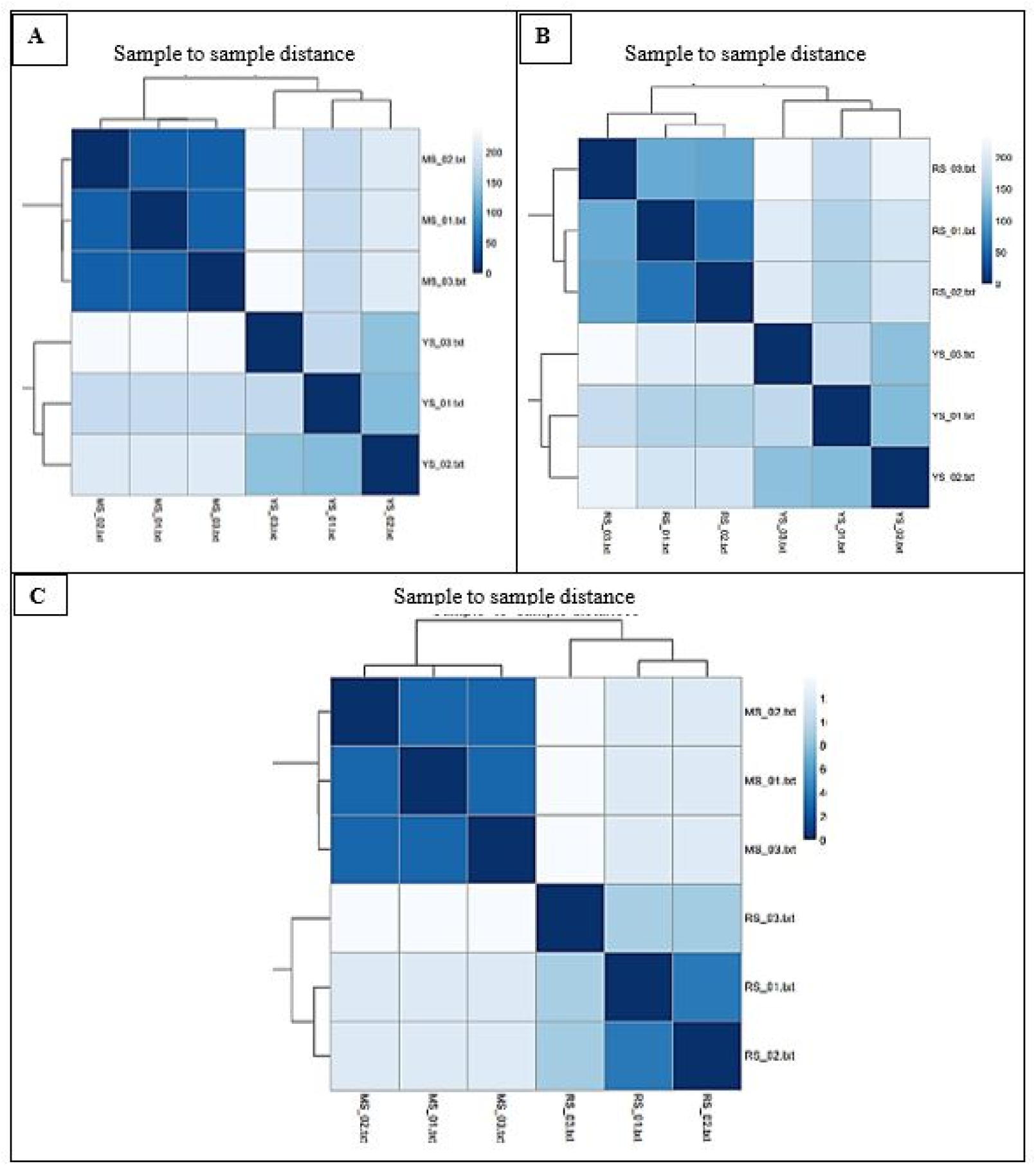
Heatmap of the sample-to-samples distances using transformed data. A. Young Stage vs Mature Stage (YS/MS), B. Young Stage vs Ripening Stage (YS/RS), and C. Mature Stage vs Ripening Stage (MS/RS)

When the young stage duplicates are compared, there are fewer similarities between them. The duplicates in the early stages, however, were grouped together and well suited for DE analysis. The relationship between gene expression at the young and ripening stages is shown in Figure 1B. The sample replicates from the ripening stage have a high degree of similarity and are grouped. Replicates from the early stages have fewer similarities but are still grouped.

Figure 1C displays the correlation of gene expression between the mature and ripening stage. There were highly similar sample replicates from the mature and ripening stage in their group in this correlation. All blocks in light blue indicate the dissimilarities of the combination of groups. All samples were suitable to proceed with the differential analysis.

### Mapping reads with Top Hat, Transcript Assembly using Cufflink, abundance estimation, comparison to reference annotation, and differential analysis

Cufflinks packages include Cuffcompare, Cuffmerge, and CuffDiff. Cufflink assembles transcripts, calculates their abundance, and tests in RNA-Seq samples for differential expression and function. It recognizes reads and assembles aligned RNA-Seq into a parsimonious collection of transcripts. Based on how many reads support each one, Cufflinks estimates these transcripts’ relative abundances (Blankenberg et al., 2010). CuffMerge was run to merge two or more transcript assemblies or the assembled transcripts from control and experimental samples and the output is the transcriptome.

Cufflinks also include Cuffdiff, which accepts the reads assembled from two or more biological conditions and analyses their differential expression of genes and transcripts, thus aiding in the investigation of their transcriptional and post-transcriptional regulation under different conditions. Data analysis was carried out using CuffDiff to obtain the list of differentially expressed genes. Cuffdiff finds significant changes in transcript expression, splicing, and promoter use. The results have reported the list of differentially expressed genes.

Gene and transcript expression analyses were studied statistically. The up-regulated and down-regulated genes and transcripts were ranked and determined using the q value and fold change. FPKM (Fragments per kilobase of transcript per million mapped fragments) value was used to evaluate the abundance of the genes in the differential group. If the absolute value of the expression log2fold change is greater than 1.5 and the false discovery rate (FDR)-corrected p-value is less than 0.05, the differential gene expression is statistically significant. A series of analyses, such as the number of expressed genes/transcripts, the number of up and down-regulated genes/transcripts and the number of differentially expressed genes/transcripts, were determined. Plots, histograms, graphs, figures and tables were constructed and studied.

Figure 2 shows the schematic diagram of the entire study workflow. To check the quality, the input dataset was uploaded to FastQC. Then, to check the quality after trimming, we will continue running trimmomatic and fast QC. After pre-processing and trimming, TopHat was used to align the reads to the Durian Reference Genome. TopHat reads RNA-Seq and is a rapid splice junction mapper. In order to locate splice junctions between exons, it first aligns the RNA-Seq readings and then analyses the mapping results. The transcripts that corresponded to the reference genome were assembled using Cufflink. Cuffdiff was then used to analyze differential expression. Additionally, mapping findings were viewed locally using the Integrative Genomics Viewer (IGV).

**Figure 2.**
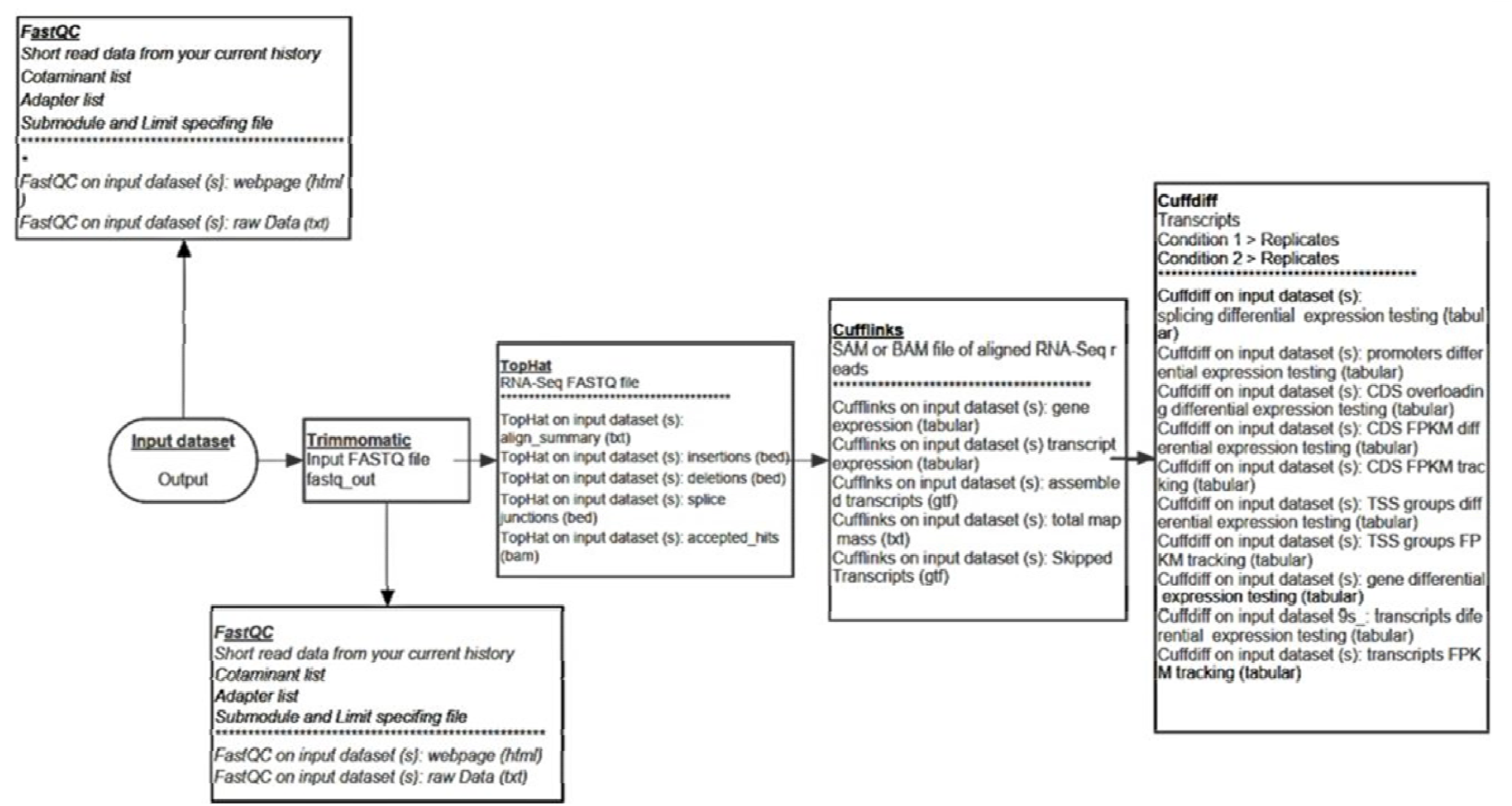
The schematic diagram showing Galaxy Workflow for CuffDif

### Comparison against reference database using Cuffcompare and novel transcript detection

The Cuffcompare program from Cufflinks was used to compare assembled transcript assemblies to annotated transcripts, which reveals novel exons, introns, and loci. Using genomic coordinates, this program generates a set of transcripts that may be used to identify both novel and well-known transcripts from the chosen reference annotation. The input for cuffcompare is cufflink GTF output and the D. zibethinus reference annotation. The output of cuffcompare is transcript accuracy files and transcript combined files that report various statistics with sensitivity and specificity values related to the accuracy of transcripts in each sample compared with reference annotation data. Cuffcompare reports a GTF file containing the “union” of all transfrags in each sample. If a transfrag is in both samples, it is only reported once in the combined gtf.

The differential expression analysis identified 280 unknown transcripts (37 and 79 up-regulated and 114 and 50 down-regulated genes in YS/MS and YS/RS) (Table S1). These sequences did not align with the Durian reference genome and might be considered unknown transcripts. These significant novel transcripts may involve different pathways or molecular networks.

Out of 280 identified novel transcripts, 183 hits the Blastn for homology search (16 and 63 up-regulated genes and 83 and 21 are down-regulated genes in YS/MS and YS/RS and were considered a novel transcript discovered in Durio zibethinus. BLASTn was used to compare the unknown sequences to the known annotated sequences available in the NCBI Genebank database (The National Center for Biotechnology Information). All the possible novel transcripts were viewed and further downloaded using IGV (Integrated Genome Viewer) for nucleotide search. The FASTA sequences of unknown transcripts were uploaded and used as a query for a BLAST search in NCBI. From the BLASTn search result, the transcripts that were annotated to the known protein-coding gene of other organisms were identified.

An unknown transcript of 97 (21 and 16 up-regulated genes and 31 and 29 down-regulated genes in YS/MS and YS/RS) did not hit any Blastn and were subjected to further analysis to search for protein domains. The transcripts with no hits in BLAST and no IPS match were shortlisted and further analyzed using Rfam and ORF. Rfam database was used to characterize sequences with no similarity to any proteins and identify homologues of known non-coding RNA families. ORF finder is a program available on the NCBI website. The program searches for the open reading frame (ORF) for all possible protein-coding regions in the sequences. Figure 3 illustrated the workflow analysis of the novel transcript.

**Figure 3.**
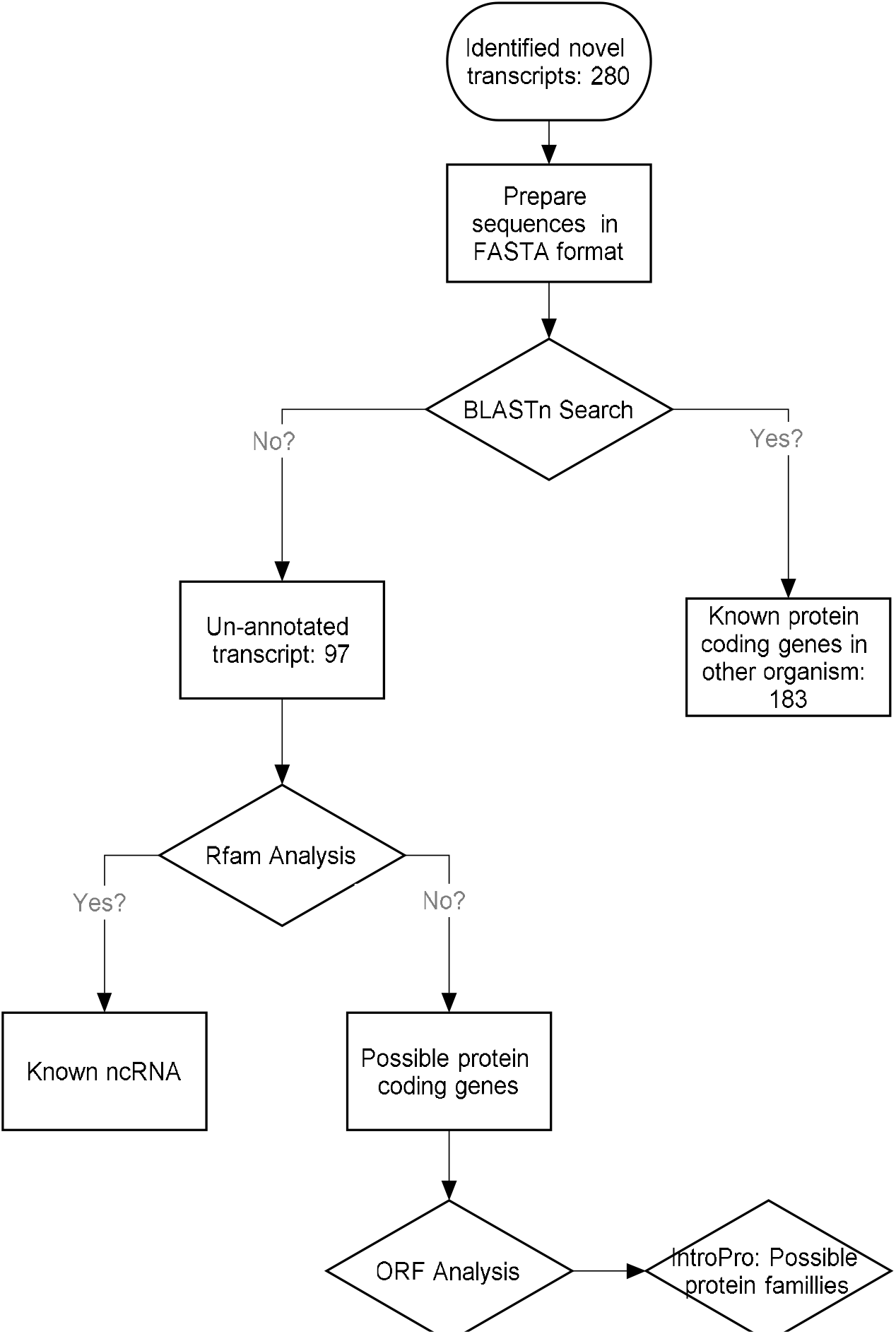
Workflow of analysis of novel transcript in durian transcriptomics.

After BLAST analysis, the public EMBL-EDI InterPro web-service database was used to analyze the unknown transcript sequences (with no hits in BLASTn) against InterPro’s signatures. The unknown sequence information (FASTA) was uploaded to the InterPro server to initiate the InterPro scan. Nucleotides were translated to amino acids automatically in the system. InterPro scan is a collection of protein families, domains and signals of peptides. Selected databases available in the analysis were identified from PHOBIUS, TMHMM, GENE3D, PROSITE_PATTERNS, SUPERFAMILY, PANTHER.

## Results

### Deep sequencing of *Durio zibethinus* transcriptome

A total of nine samples collected from three biological replicates of *D. zibethinus* from different growth stages were subjected to RNA-Seq using the tuxedo package. The paired-end reads’ sequence was aligned to the *Durio zibethinus* reference genome using TopHat (Galaxy Version 2.1.1). This tool accepts files in Sanger FASTQ format. TopHat has produced four output files namely; junctions (BED track of junction), insertions, deletions and accepted_hits (BAM format of alignment files). TopHat default settings were used: 20 for a maximum number of alignments to be allowed, 2 for final read mismatches and yes to the output of unmapped reads in BAM format. The tool has been chosen as a mapping tool because it can create a database of splice junctions based on the model gene’s annotation (Trapnell et *al*. 2012). Table 1 below showed the mapping results in the 84.5-90.6% range.

**Table 1.**
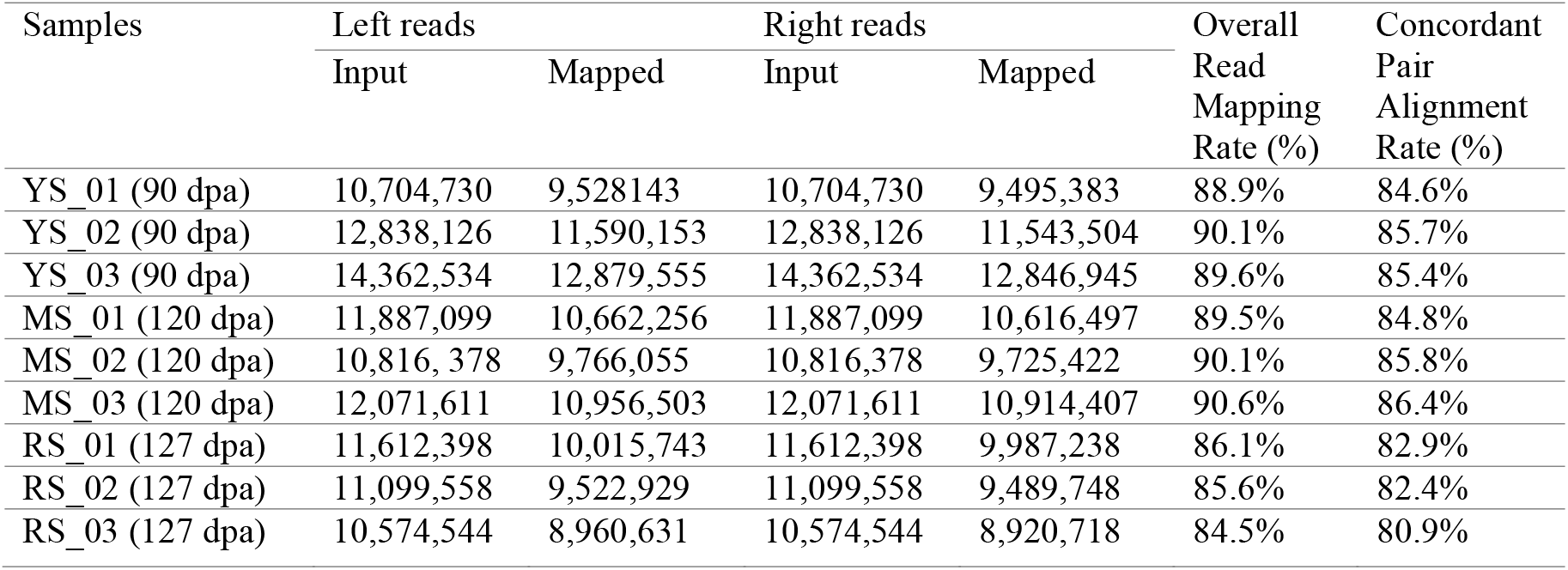
Mapping results against *Durio zibethinus* Reference Genome (Reference Guided Assembly) using TopHat Galaxy ver. 2.1.1

Cufflink was used to assemble the transcripts, estimate their abundance using the FPKM value and perform the differential expression testing. Cufflinks were carried out to reconstruct the transcripts together with Tophat2. This exon-guided assembly helped capture the differentially expressed genes in samples with well-annotated genomes. A total of 110 million high-quality reads were analyzed for all nine samples, individually identifying 76,700-89,117 transcripts and 561,211-646,292 exons after the mapping process.

To obtain the unannotated transcripts, the known transcripts were filtered out with Cuffcompare. Cuffcompare was used to compare the cufflink assemblies to reference annotation files. To identify new transcripts from known ones, all of the reconstructed transcripts were compared individually by aligning to the annotation of the durian reference genome. The total query mRNAs is 88,000 in 44,509 loci (78,791 multi-exon transcripts); (15,910 multi-transcript loci, ∼2.0 transcripts per locus. The total number of reference mRNAs is 76,695 in 43,808 loci (69,546 multi-exon). The total super-loci w/ reference transcripts are 42,768. The novel exons, introns and loci were identified in the samples of young stage. The results as follows: novel exons: 3,438/313,476 (1.1%), novel introns: 1,657/245,129 (0.7%), and novel loci: 1,197/44,509 (2.7%). The missed exons is 9/295602 (0.0%), missed introns is 282/237,421 (0.1%) and missed loci is 5/43808 (0.0%). These novel exons’ existence was shown in the alignment viewed using Integrated Genome Viewer (IGV) and differential expression testing results.

### Differential Gene Expression (GE) analysis with CuffDiff

Differentially expressed genes and transcripts were considered statistically significant if the absolute value of the expression log2 fold change > 1.5 and < -1.5. A significantly DEG selection was also performed by filtering the genes with FDR-corrected p-value, q value <0.05.

The outputs from Cuffdiff were as follows, Splicing differential expression testing, Promoter differential expression testing, CDS overloading differential expression testing, CDS FPKM differential expression testing, CDS FPKM tracking, TSS groups differential expression testing, TSS groups FPKM tracking, Gene differential expression testing, Gene FPKM tracking, Transcript differential testing and Transcript FPKM tracking.

After the readings were mapped to the reference genome with TopHat, Cufflink was used to put together the transcripts. CuffDiff was used to analyze the expression levels of differentially expressed genes (DEGs) at three different stages. Only genes with significant expression with q value 0.05 (5% of a gene for each comparative group, false discovery rate, FDR) were considered. The DEGs were compared across three stages of development. The total number of up-regulated and down-regulated expressed genes was listed in Table 2. During the transition from a young to a mature stage, the number of down-regulated genes was significantly higher than that of up-regulated genes. On the other hand, the number of up-regulated genes was higher than the number of down-regulated genes in the young to ripening stages. During the transition from the young to the mature stage, the expression of 1,496 up-regulated genes and 3,738 down-regulated genes were analyzed further. During the transition from the mature to the ripening stage, the expression of 2,153 genes and 926 genes was upregulated and downregulated.

**Table 2.** Gene Differential Expression for three compared groups using CuffDiff.

| Type | YS/MS<br>(Annotated gene) | YS/RS<br>(Annotated Genes) | MS/RS<br>(Annotated Genes) |
| --- | --- | --- | --- |
| Up-regulated genes | 1,496 | 2,154 | 2,153 |
| Down-regulated genes | 3,738 | 927 | 926 |

Table 3 shows the top 10 up and downregulated among the differentials expressed genes of YS/MS and MS/RS.

**Table 3.** Top 10 up and down-regulated among the differentially expressed genes

| Gene ID | Gene name | log 2 (fold change) | q_value | Stage | Up/Down |
| --- | --- | --- | --- | --- | --- |
| LOC111304630 | LOB domain-containing protein 18, (Lateral Organ Boundaries) | 9.1229 | 0.000312596 | YS/MS | Up |
| LOC111311552 | polygalacturonase inhibitor-like | 8.43397 | 0.000312596 | YS/MS | Up |
| XLOC_006973 | novel transcript | 8.17641 | 0.000312596 | YS/MS | Up |
| LOC111292650 | triacylglycerol lipase 2-like | 7.55833 | 0.000312596 | YS/MS | Up |
| LOC111293555 | benzyl alcohol O-benzoyltransferase-like, transcript variant X1 | 7.10972 | 0.000312596 | YS/MS | Up |
| LOC111307587 | *uncharacterized LOC111307587* | 6.58076 | 0.00129227 | YS/MS | Up |
| LOC111282509 | *uncharacterized LOC111282509* | 6.55632 | 0.00343537 | YS/MS | Up |
| LOC111281105 | protein DETOXIFICATION 40-like | 6.45873 | 0.000312596 | YS/MS | Up |
| LOC111318443 | bidirectional sugar transporter N3 | 6.3118 | 0.000312596 | YS/MS | Up |
| LOC111315580 | synaptonemal complex protein 1-like | 6.25794 | 0.000312596 | YS/MS | Up |
| LOC111278395 | endoglucanase-like | -11.0334 | 0.000833693 | YS/MS | Down |
| LOC111315188 | allene oxide synthase 1, chloroplastic-like | -10.662 | 0.0496018 | YS/MS | Down |
| LOC111291309 | abscisate beta-glucosyltransferase-like | -10.2951 | 0.000312596 | YS/MS | Down |
| LOC111305464 | auxin-induced in root cultures protein 12-like | -9.74314 | 0.0108235 | YS/MS | Down |
| LOC111295419 | protein CDI-like | -9.7041 | 0.00999511 | YS/MS | Down |
| LOC111288324 | UDP-glucuronate 4-epimerase 6-like | -9.65161 | 0.00451342 | YS/MS | Down |
| LOC111281364 | dehydration-responsive element-binding protein 1A-like | -9.37327 | 0.000582785 | YS/MS | Down |
| LOC111303526 | heavy metal-associated | -9.28425 | 0.000312596 | YS/MS | Down |
| LOC111312815 | chaperone protein dnaJ 11, chloroplastic-like | -9.25731 | 0.038914 | YS/MS | Down |
| LOC111284909 | asparagine synthetase [glutamine-hydrolyzing] 1 | -9.21616 | 0.00343537 | YS/MS | Down |
| LOC111317014 | probable mannitol dehydrogenase | 9.51845 | 0.0167276 | MS/RS | Up |
| LOC111311521 | polygalacturonase-like | 9.48951 | 0.00020799 | MS/RS | Up |
| LOC111278395 | endoglucanase-like | 9.03817 | 0.00206807 | MS/RS | Up |
| LOC111295536 | exocyst complex component EXO70H1-like | 8.82363 | 0.00516878 | MS/RS | Up |
| LOC111287629 | small heat shock protein, chloroplastic-like | 8.76183 | 0.00727459 | MS/RS | Up |
| LOC111300662 | stress enhanced protein 2, chloroplastic-like | 8.34376 | 0.0435014 | MS/RS | Up |
| LOC111291309 | abscisate beta-glucosyltransferase-like | 7.96966 | 0.00020799 | MS/RS | Up |
| LOC111278572 | inorganic pyrophosphatase 2-like | 7.82673 | 0.00020799 | MS/RS | Up |
| LOC111311985 | glutathione S-transferase U8-like | 7.71176 | 0.00020799 | MS/RS | Up |
| LOC111285272 | *uncharacterized LOC111285272* | 7.67617 | 0.00287618 | MS/RS | Up |
| LOC111311552 | polygalacturonase inhibitor-like | -8.47809 | 0.000208 | MS/RS | Down |
| LOC111294291 | *uncharacterized LOC111294291* | -7.81045 | 0.000208 | MS/RS | Down |
| LOC111311438 | transport and Golgi organization 2 homolog | -7.73797 | 0.0112713 | MS/RS | Down |
| LOC111282412 | LOB domain-containing protein 42-like | -6.42836 | 0.000208 | MS/RS | Down |
| LOC111283878 | *uncharacterized LOC111283878* | -6.31947 | 0.000208 | MS/RS | Down |
| LOC111313342 | myb-related protein 308-like | -6.15936 | 0.0211934 | MS/RS | Down |
| LOC111297709 | *uncharacterized LOC111297709* | -6.06598 | 0.0431503 | MS/RS | Down |
| LOC111277115 | *uncharacterized LOC111277115* | -6.04314 | 0.000208 | MS/RS | Down |
| LOC111297280 | fructose-bisphosphate aldolase 1, chloroplastic-like | -5.95271 | 0.000208 | MS/RS | Down |
| LOC111299239 | solute carrier family 25 member 44-like | -5.92814 | 0.000208 | MS/RS | Down |

### Hierarchical clustering of gene expression profiles across the three growth stages using RPKM value

All up-regulated DEGs were clustered to examine gene expression patterns and identify the significant gene cluster in each studied differential group. Figure 4 displays the heatmap analysis of complete DEGs in three hierarchical growth phases. Each distinct group’s gene expression patterns can be seen in this heatmap. Blue and orange denote up and down-regulated genes at the young, mature, and ripening stages, respectively. The rows reflect measurements from various genes, whereas the columns represent multiple samples. Orange typically denotes a gene with low expression, while blue denotes a gene with high expression. Based on similarity, hierarchical clustering arranges the rows and/or columns. A side-by-side dendrogram indicating gene grouping is displayed, and a sample is displayed at the bottom of the heatmap.

**Figure 4.**
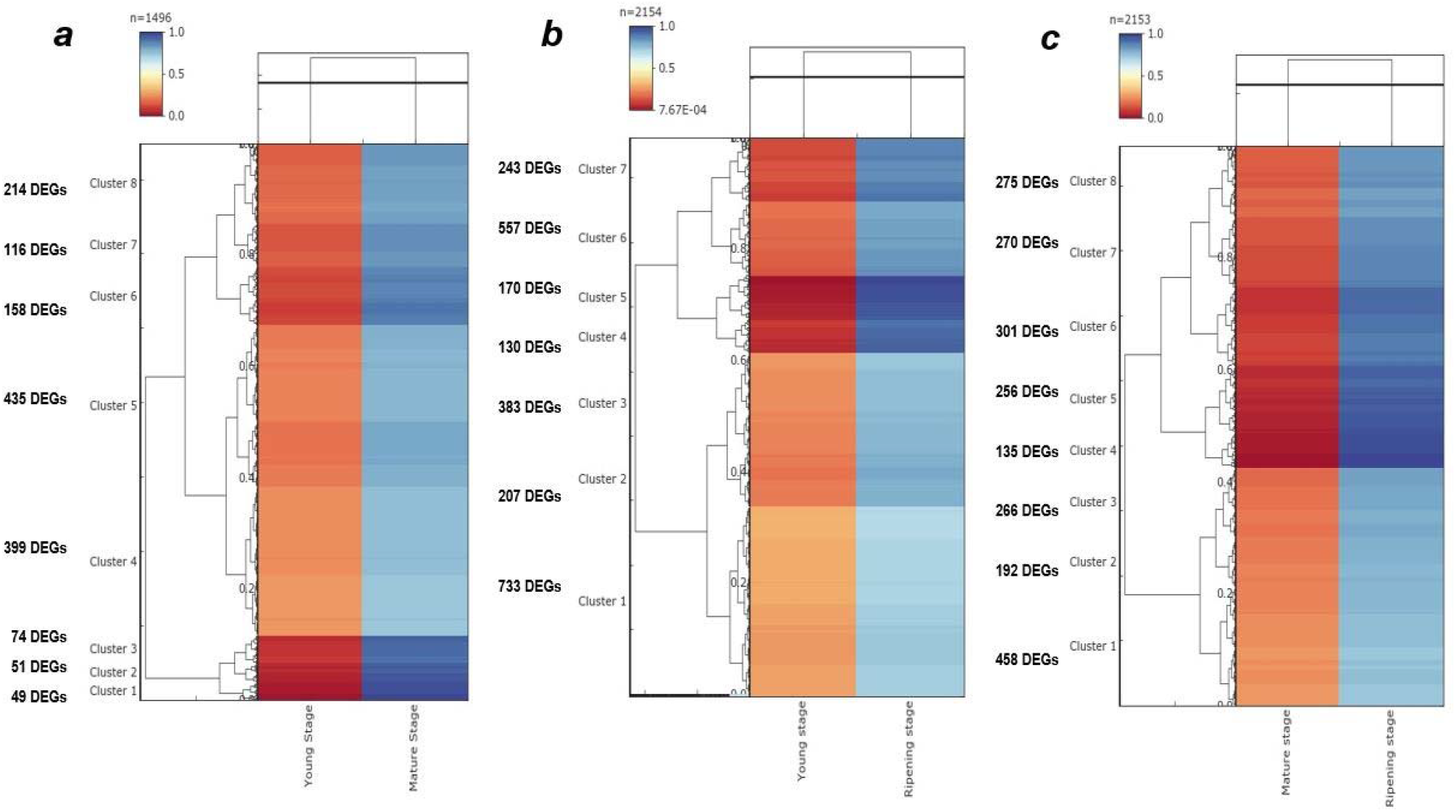
Heatmap analyses hierarchical clustering of whole DEGs in the a. up-regulated transition between the young and mature stage, b. up-regulated in transition between young and ripening stage and c. up-regulated in transition between mature and ripening stage. Clusters were generated using a hierarchical-means method based on 1,496 DEGs. Note that data using RPKM value after normalization with feature scaling. Each column represents a test sample. Each row represents a change of gene expression of different colours, blue bands indicate up-regulation, and orange bands

Eight (YS/MS), seven (YS/RS), and eight (MS/RS) main clusters were identified by hierarchical clustering and were denoted by coloured bars between the dendrogram and heatmap. To analyze significant expression differences, the FDR of 0.05 and the maximum value of log2 (ratio of young/mature/ripening)| 1 were chosen as cut-offs.

Genes with comparable expression patterns were grouped during the developmental and ripening stages of growth, and the patterns could be accessed through colour variations. An obvious comparison could be made between all the different levels of the expression pattern.

The hierarchical clustering for 1,496 up-regulated DEGs in the transition from the young to mature stage was further discussed. There were all grouped into eight clusters: Cluster 1 (49 DEGs), Cluster 2 (51 DEGs), Cluster 3 (74 DEGs), Cluster 4 (399 DEGs), Cluster 5 (435 DEGs), Cluster 6 (158 DEGs), Cluster 7 (116 DEGs) and Cluster 8 (214 DEGs). Of the 8 clusters, 3 clusters (clusters 1, 2 and 3) had the most important DEGs released and associated with response to stress, cuticle formation, receptor and protein kinases, organ development, general transporter, transcription factor, sugar transporter, hormone signalling, starch synthesis, ROS generation, DNA repair, and cell wall degradation.

The hierarchical clustering for 2,154 up-regulated DEGs in the transition from young to ripening stage was further discussed. There were all grouped into seven clusters: Cluster 1 (733 DEGs), Cluster 2 (207 DEGs), Cluster 3 (383 DEGs), Cluster 4 (130 DEGs), Cluster 5 (170 DEGs), Cluster 6 (557 DEGs), and Cluster 7 (243 DEGs). Of the 7 clusters, 2 clusters (clusters 4 and 5) had the most important DEGs expressed. The results showed that the cluster with the majority of genes with an increased expression upon transition from young to ripening stage was associated with response to stress, transcription factor, general transporter, sulphate transporter, and sugar transporter.

The hierarchical clustering for 2,153 up-regulated DEGs in the transition from mature to ripening stage was further discussed. There were all grouped into eight clusters: Cluster 1 (458 DEGs), Cluster 2 (192 DEGs), Cluster 3 (266 DEGs), Cluster 4 (135 DEGs), Cluster 5 (256 DEGs), Cluster 6 (301 DEGs), Cluster 7 (270 DEGs) and Cluster 8 (275 DEGs). Of the 8 clusters, 2 clusters (clusters 4 and 5) had the most significant DEGs expressed. The results showed that the cluster with the majority of genes with an increased expression upon transition from the mature to the ripening stage was mainly involved in response to stress, cuticle formation, cell wall degradation, hormone signalling, Ca2 signalling, general transporter, and receptor and protein kinases.

### Nucleotide search using BLASTn for 280 unknown transcripts

#### Annotation of novel transcripts

BLASTn was used to compare the unknown sequences to the known annotated sequences available in the NCBI Genebank database (The National Center for Biotechnology Information). All the possible novel transcripts were viewed and further downloaded using IGV (Integrated Genome Viewer) for nucleotide search. The FASTA sequences of unknown transcripts were uploaded and used as queries for a BLAST search in NCBI. From the BLASTn search result, the transcripts that were annotated to the known protein-coding gene of other organisms were identified.

E-value sorts of blast results by default (best hit in the first line). The smaller the E-value, the better the match. Blast hits with an E-value smaller than 1e-50 include database matches of very high quality. Blast hits with an E-value smaller than 0.01 can still be considered a good hit for homology matches. In other words, the lower the E-value or, the closer it is to zero, meaning the more significant the hits (Syngai et *al*., 2013). In this study, the expected threshold is an E-value <1.0E-3 with a filter of low complexity regions to avoid misinterpretation of the results.

There were identified 280 unknown transcripts with multiple and single exon structures, extracted from DEGs analysis, that did not annotate with the Durian reference genome and might be considered as an unknown significant transcript as illustrated in Figure 5. The size of new unknown transcripts had distributed between 150 bp and 8 kb. These unknown transcripts were searched through blast2GO (blastn) to see whether they could be matching to any taxa other than *D. zibethinus* and to investigate their possible novelties. Out of 280 identified novel transcripts, 183 hits the Blastn for homology search (16 and 63 up-regulated genes and 83 and 21 are down-regulated genes in YS/MS and YS/RS). 183 unknown transcripts were considered a novel transcript discovered in *Durio zibethinus*, while 97 were subjected to further analysis to search for protein domains.

**Figure 5:**
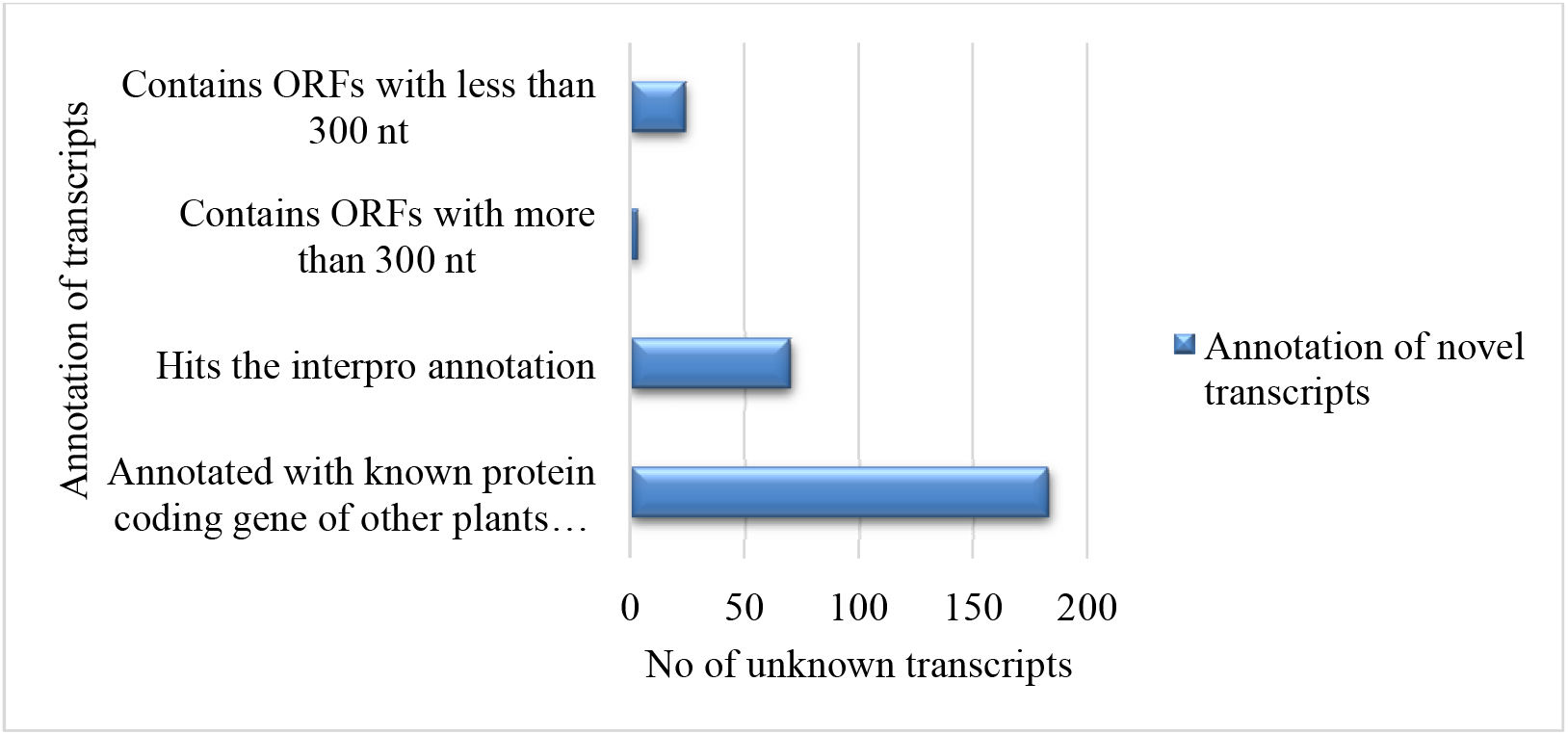
Annotation of unknown transcripts.

A large number of transcripts that did not show any hits to any of the databases but were significantly and differentially regulated were identified in this study. These unknown transcripts may involve different pathways or molecular networks during young, mature, and ripening stages. The novel transcript shows an identical or homologous structure of cDNAs of other species, which is absent from the Durian reference genome. Novel exon is an exon present in known or unknown transcripts and not supported by the Durian reference genome. Novel locus is a contiguous stretch of the genomic region from which the above transcript and exons are expressed. This research solely focused on the transcript that indicated up-regulated of differentially expressed groups only.

Further differential expression analysis revealed 37 unknown transcripts in the up-regulated and 131 unknown transcripts in the down-regulated transition from the young to the mature stage. In the transition of young to the ripening stage, there were 79 unknown transcripts in the up-regulated and 50 unknown transcripts in the down-regulated. Since the priority of the study is to recognize only significant transcripts, only differentially expressed of unknown transcripts were filtered for further analysis.

#### Analysis of protein domains using InterPro Scan

An unknown transcript of 97 (21 and 16 up-regulated genes and 31 and 29 down-regulated genes in YS/MS and YS/RS) did not hit any Blastn. These 97 novel transcripts did not show any similarity with any sequences available in the gene bank’s nucleotide database. The public EMBL-EDI InterPro web-service database was used to analyze the 97 unknown transcript sequences (with no hits in BLASTn) against InterPro’s signatures. The unknown sequence information (FASTA) were uploaded to InterPro server to initiate the InterPro scan. Nucleotides were translated to amino acids automatically in the system. As a result, 70 unknown transcripts were hits the Interpro annotation, and the other 27 unknown transcripts were categorized under “No IPS match”.

### Analysis of Rfam and ORF of unknown sequences with no IPS match

#### Non-coding RNA analysis using the rfam database

Rfam database was used to characterize sequences with no similarity to any proteins and identify homologues of known non-coding RNA families. The RNA families were classified into three functional classes: non-coding RNA genes, structured cis-regulatory elements and self-splicing RNAs. Rfam database is represented by multiple sequence alignments, consensus secondary structures and covariance models (Cms). The transcripts with no hits in BLAST and no IPS match were shortlisted and further analyzed using Rfam. From the results of the Rfam search, the transcripts that are having hits are as known non-coding genes. Non-coding RNAs analysis can be carried out by constructing the co-expression network to characterize the functions of non-coding RNAs (Kalvari et *al*., 2018). The analysis of unknown sequences displayed no hits results in the Rfam database, indicating that all unknown sequences are potentially protein-coding or small peptides.

#### ORF Prediction results

ORF finder is a program available on the NCBI website. The program searches for the open reading frame (ORF) for all possible protein-coding regions in the sequences. Any ORF and its protein translation are returned by the software. The six horizontal bars corresponding to one of the possible reading frames with each direction of the DNA would be possible for three reading frames. The amino acid results were used to search for various protein databases using BlastP or Blastx for finding similar sequences. The longest ORF (usually >100 codons) from the methionine start codon was a good prediction of protein-encoding sequence because it contains rich features suitable for most gene prediction programs (Haoyu et *al*., 2008). Generally, ORF lengths of more than 300 nucleotides or at least 100 codons were considered for proteome analysis. ORFs lengths with less than 100 codons were classified under small ORFs. Previous research conducted revealed that small ORFs were also functional and significant peptides. These peptides are also involved in gene expression and contribute to the role of any biological pathways (Yazhini, 2018).

## Discussion

This study presents transcriptome analysis by deep RNA sequencing to study durian genetic expression in their three stages of growth. The study reveals the scope of comprehensive RNA Seq in Durian to find novel transcripts as annotation of the *D. zibethinus* genome is incomplete. We found several known, unknown transcripts and exons with a significant differential expression that may play a role in the development of Durian growth.

It is crucial to map cDNA reads to their genomic regions to identify novel transcripts and measure accurate expression based on RNA-seq. We used TopHat to map cDNA reads to reference the genome, as it is designed to work with splice site detection. As a result of the mapping, approximately 90% of reads were mapped uniquely, which indicates a high quality of mapping and reliability of gene structures predicted from such mapping. The significant difference in the number of up-and-down-regulated genes between young and ripening stages may be due to increased bioactivity in the late development stage. During the transition from mature to ripening, the phenomenon with a higher number of up-regulated genes is associated with a few biochemical changes that may occur at the ripening stage to help fruit development. For a broader range of analysis, the outcome from Cuffdiff was used for further study of whole DEGs from three stages of growth. CuffDiff was chosen due to its high mapping rate and highly known and annotated gene in the top significant DEGs.

A comprehensive study from the heatmap indicates that response to stress, hormone signalling, cuticle formation, general transporter, receptor and protein kinase and transcription factors are highly regulated gene functions for all growth stages. A significant increase in gene number represents transcription factor (TF) from the developmental to ripening stage. All growth stages highly expressed genes related to hormone signalling, ethylene signalling (ACO, ACS), and plant hormone signalling (Auxin, ABA). MGL gene isoforms were found in LOC111308047 (methionine gamma-lyase-like) and LOC111290112 (methionine gamma-lyase-like) expressed in the development and ripening stages, respectively. E3 ubiquition-protein ligase is highly expressed in all stages of growth to support multiple plant development. Zinc finger protein is also highly expressed in all stages of growth for stress response and defence. Xyloglucan endotransglucosylase/ hydrolase is highly expressed to promote the durian fruit softening in the late growth stage.

Our study focuses on the 280 unknown transcripts for further analysis. 183 unknown transcripts were annotated with known protein-coding genes of other plant species. It is considered novel related to first found in *D. zibethinus*. 70 unknown transcripts with no Blastn hit matched the identifier from Interpro annotation. 27 unknown transcripts were categorized under “No IPS match”. Out of 27, 3 unknown transcripts with ORFs sequences with more than 300 nt were identified and 24 with less than 300 nt were considered small ORFs.

In the analysis, 183 unknown transcripts were annotated to the known protein-coding gene of other plant species closely related to the Durian genome. These annotated transcripts were known as novel related, which was first discovered in durian. Top blastn results were shown a homology (80-100%) to *Bombax ceiba, Gossypium hirsutum, Gossypium raimondii, Herrania umbratica, Juglans regia, Theobroma cacao* and many other plant species were indicated further in the list. The top BLASTn results as presented in Figure 6 were matched with previous results that claimed *Theobroma cacao* (cacao; used in chocolate) and members of the *Gossypium* genus (cotton) are the relative genome that’s available include only the more distantly related cash crops within the more prominent Malvaceae family (Teh et *al*., 2017). Malvaceae or the mallows is a family of flowering estimated to contain 244 genera with 4,225 known species, with known members of economic importance including okra, cotton, cacao and durian. The Malvaceae s.l. (hereafter merely “Malvaceae”) comprise nine subfamilies: Bombacoideae, Brownlowioideae, Byttnerioideae, Dombeyoideae, Grewioideae, Helicteroideae, Malvoideae, Sterculioideae, Tilioideae.

**Figure 6:**
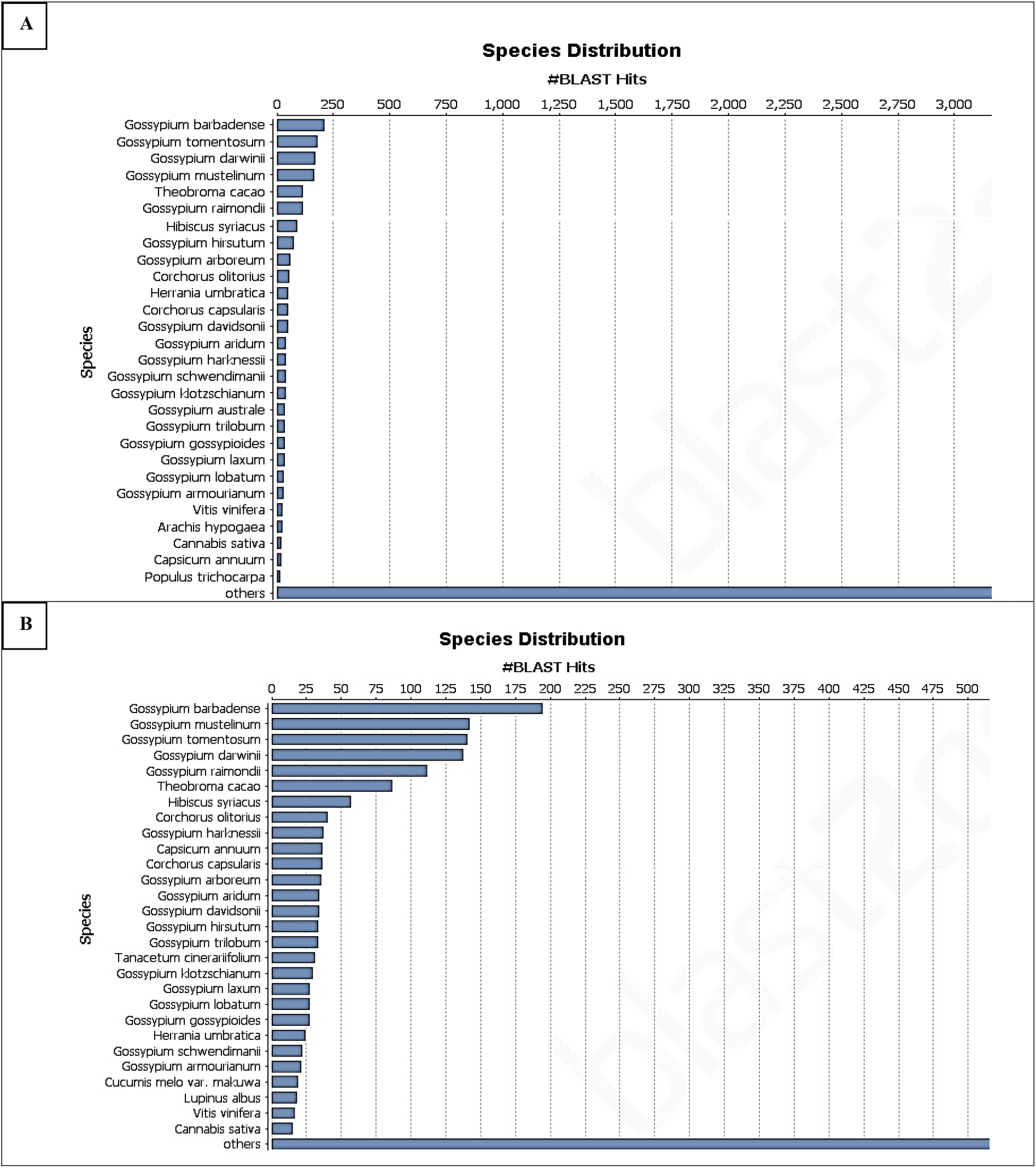
Distribution of BLAST results for unknown transcripts by species is shown as the percentage of the total; homologous sequences (with an E-value <1.0E-3). All plant proteins in the NCBI nr database were used for homology search and the best hit of each sequence was used for analysis. (A) YS/MS, (B) YS/RS.

**Figure 7:**
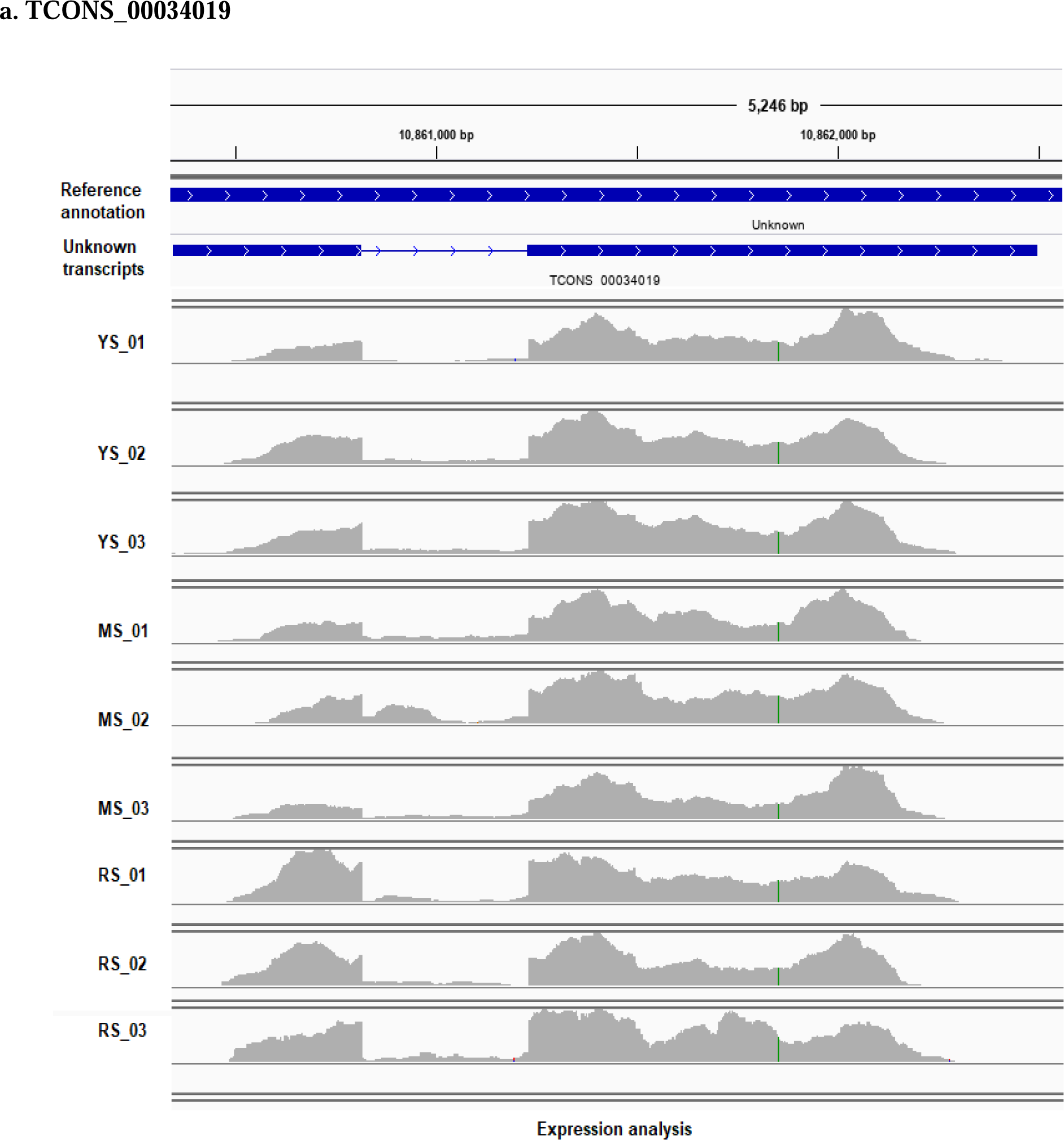
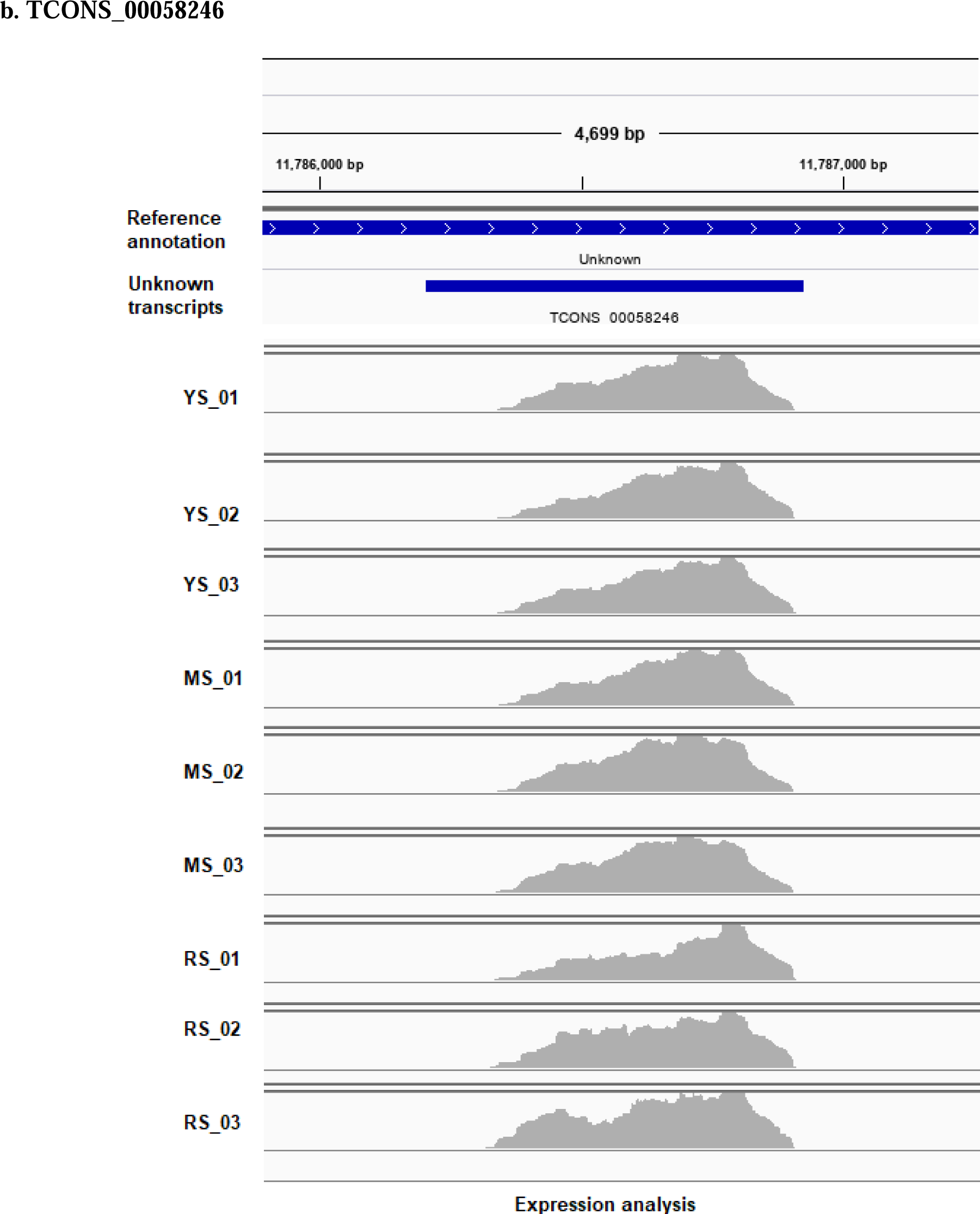
Expression analysis of unknown transcript that hits GO terms using IGV. (a) Transcript id of TCONS_00034019 and (b) Transcript id of TCONS_00058246.

Under the category of unknown transcripts with no Blastn hit, two unknown transcripts hits GO terms; TCONS_00034019 and TCONS_00058246. TCONS_00034019 [locus position: NW_019167937.1:10860347-10862497], hits the InterPro GO Names for Biological Process (P) P:GO:0046622; P: positive regulation of organ growth. The sequences were extracted from IGV and uploaded to BLASTx to see the match. The top hits for the sequences were identified as the auxin-regulated gene involved in organ size [Theobroma cacao] with a similarity of 92.1% and E value of 3e-17 as indicated in figure 6a. TCONS_00058246 [locus position: NW_019168015.1:11786205-11786925] hits the InterPro GO Names for Molecular Function (F), namely F:GO:0005509; F: calcium ion binding. The sequences were extracted from IGV and uploaded to BLASTx to see the match. The sequences’ top hits were identified as an auxin-regulated gene involved in organ size [Theobroma cacao], with a similarity of 93% and an E-value of 3e-17. Figure 6 shows the expression analysis of an unknown transcript that hits GO terms. InterPro scan is a collection of protein families, domains and signals of peptides. The selected database in the analysis are identified from PHOBIUS, TMHMM, GENE3D, PROSITE_ PATTERNS, SUPERFAMILY, PANTHER as indicated in Figure 6b.

Under the category of sequences with no BLASTx hit and no IPS match, there were three ORFs sequences with more than 300 nt in length. The locus XLOC_004344 (consists of TCONS_00009381) with 1650 nt; 549 aa was classified as an unknown protein-coding gene. This transcript was differentially expressed between the young and mature stages in the up-regulated level and did not annotate in the reference Durian genome. Comparative sequence analysis using SMARTBLAST based on UniProtKB/Swiss-Prot detected sequence similarity with DOMAIN: Retropepsins; pepsin-like aspartate proteases uncharacterized protein LOC100803389 [*Glycine max*] with e-value of 2e-06 and identity 23.12%. The locus XLOC_035884 and XLOC_008159 (consists of TCONS_00076984 and TCONS_00017315) with 324 nt; 107 aa and 318 nt; 105 aa, were classified as unknown protein-coding genes. These transcripts were differentially expressed between the young stage and ripening stage in the downregulated level with and also did not annotate in the reference Durian genome. Comparative sequence analysis using SMARTBLAST based on UniProtKB/Swiss-Prot detected sequence similarity with DUF4492 domain-containing protein [*delta proteobacterium NaphS2*] with e-value of 0.007 and identity of 46.67% and hypothetical protein CTI12_AA006750 [*Artemisia annua*] with e-value of 9e-11 and identity of 50.0%.

One of the critical features of RNA-Seq is the identification of novel unannotated genes expressed in the transcriptome, which is impossible with any other technique. We found several novel transcripts expressed in the young and ripening stages of growth in *D. zibethinus*. Comparatively, more transcriptional loci were found in the ripening stage the young stage, indicating increased transcriptional activity in the ripening stage. For validation of novel transcripts, we have used straight forward approach consisting of cross-verification of novel transcripts in BLAST and IGV.

This research has shown that these gaps must be filled to assist future RNA-Seq studies. While RNA-Seq can detect novel gene structures without prior annotation information, rigorous detection and validation approaches are required to exclude false transcripts, particularly in the case of low coverage and short readings provided by most current NGS technologies. However, software such as Cufflinks (Trapnell et al., 2010) is available for transcript reconstruction from RNA-Seq data, which can be used for transcript assembly, given that the sequencing is done with adequate coverage and more extended readings.

## Conclusion

In conclusion, we provided RNA-seq with a sufficient depth of 150bp reads generated by the Illumina sequencing platform of durian, resulting in the discovery of novel transcripts and exons. The results assessed in this study provide detailed information on the durian transcriptome during different stages of growth. In addition, our findings have filled the gaps in durian molecular genetics.

## Acknowledgement

The author wants to acknowledge that the research funding for this project was provided by AIMST University, Malaysia under its internal research grant scheme.

## Conflict of Interest

The author declares no conflict of interests. The data have not been published and are not under consideration elsewhere. The author has approved the submission of the manuscript.

## Ethical Compliance

This paper does not contain any studies involving humans or animals.

## Supporting information

Table S1 - 280 unknown transcripts from Cuffdiff (XLS)

Table S2 - 183 unknown transcripts that hits for BLASTn (XLS) Table S3 - 97 unknown transcripts that no hits for BLASTn (XLS)

Table S4 - 70 unknown transcripts that have InterPro Scan results (XLS) Table S5 - 27 sequences with no BLASTx hits and no IPS match (XLS)

Table S6 - Results analysis of sequences with no BLASTx hit and no IPS match (XLS)

**Figure S1.**
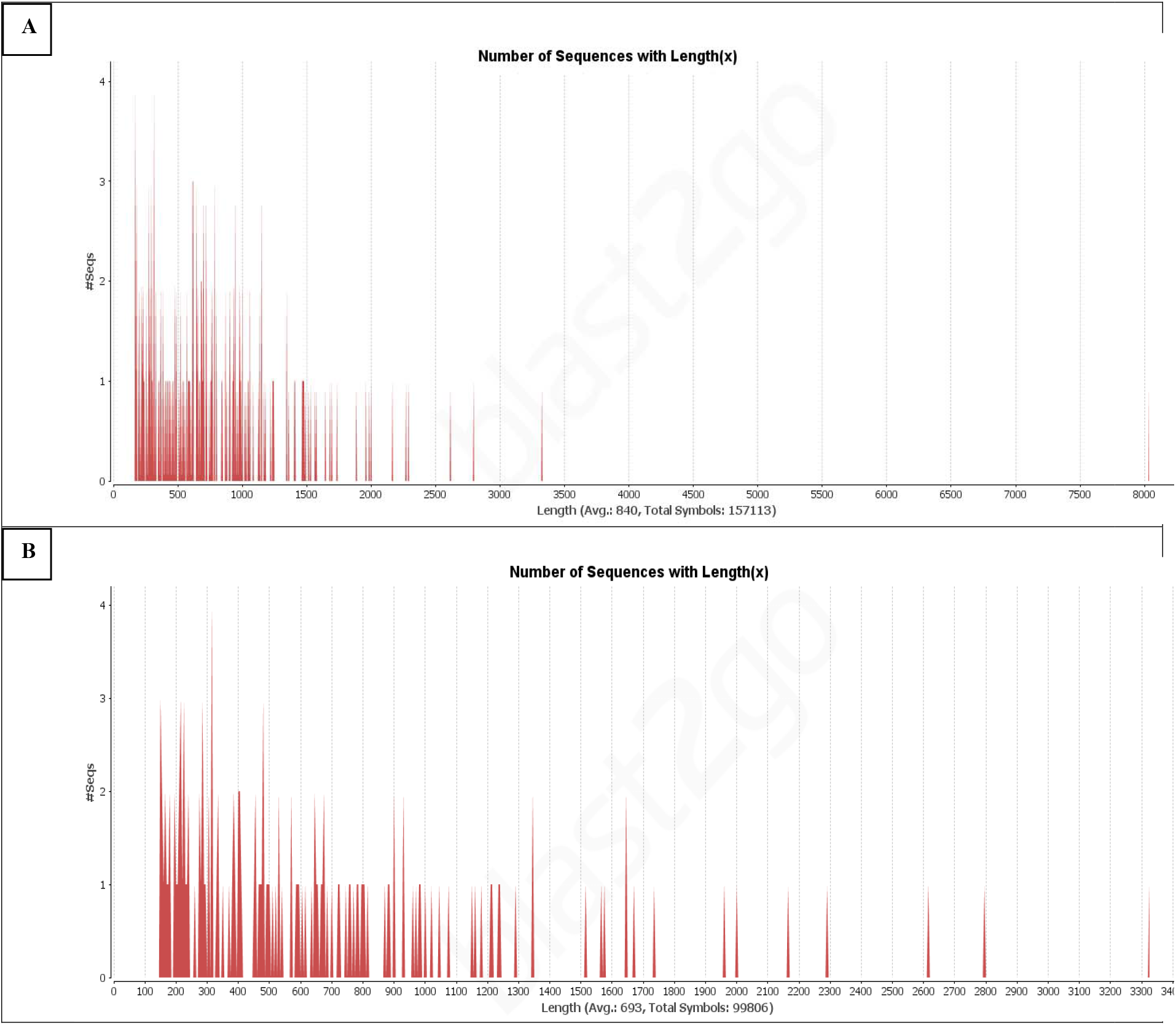
Effect of query length on the percentage of significant matches for unknown transcripts. The cut-off value was set at 1.0E-3. (A) YS/MS, (B) YS/RS

## Notes

### Competing Interest Statement

The authors have declared no competing interest.

